## Supplemental Information for "Distinct Activation Mechanisms of CXCR4 and ACKR3 Revealed by Single-Molecule Analysis of their Conformational Landscapes"

**This PDF file includes:**

Figures S1 to S10
SI References

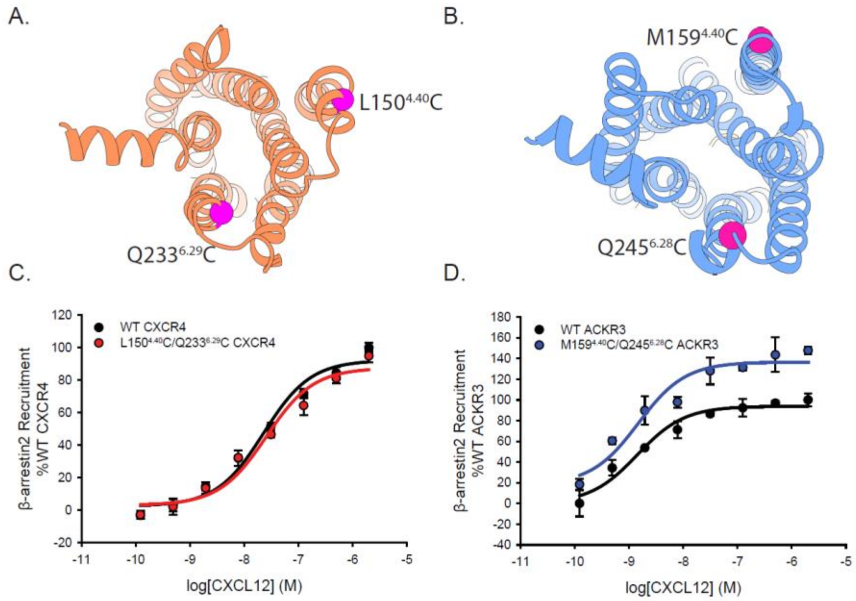


**Fig. S1.** Cysteine mutations on TM4 and TM6 of CXCR4 and ACKR3 do not impair CXCL12 mediated β-arrestin2 recruitment observed by BRET. A, B) Locations of the cysteines (magenta spheres) on CXCR4 (A) and ACKR3 (B). C, D) Dose-response BRET-based arrestin recruitment to WT receptors and Cys-containing (C) CXCR4 and (D) ACKR3. Data represents averages of three independent experiments normalized to the WT receptor recruitment measured on the same experimental plate as the mutants.

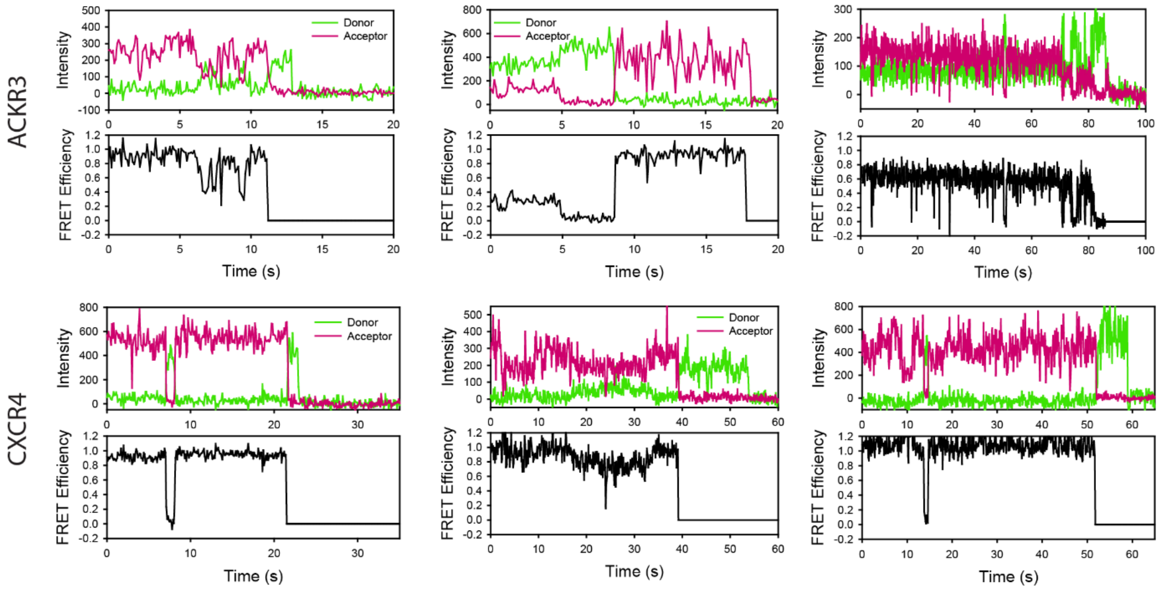


**Fig. S2.** Example single-molecule traces of apo-ACKR3 and apo-CXCR4. Donor intensity, acceptor intensity and apparent FRET efficiency time traces are colored green, magenta and black, respectively.

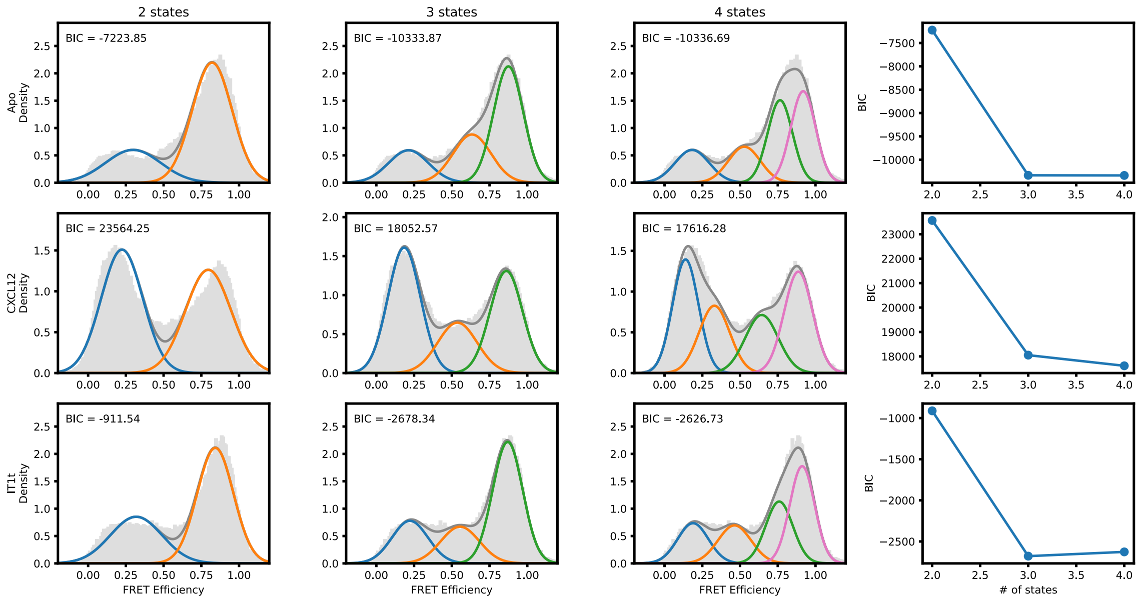


**Fig. S3.** Quantitative evaluation of the appropriate number of FRET states required to model the CXCR4 smFRET distributions. SmFRET data for CXCR4 recorded in the apo-state (first row), and in the presence of agonist $CXCL12_{WT}$ (second row) and small-molecule ligand IT1t (third row) were globally analyzed by Hidden Markov analysis assuming the presence of two, three, or four FRET states. For each condition, the overall apparent FRET efficiency histogram is shown in gray. Each distribution was fit to a Gaussian Mixture Model (GMM) using the Python package scikit-learn, and the Bayesian Information Criterion (BIC) was calculated. Individual gaussian components are shown as colored lines and the total fitted envelopes are shown as dark gray lines. The values of the BICs were used to evaluate the appropriate model complexity as described in the text. The results indicate that a three-state model is sufficient to describe the data.

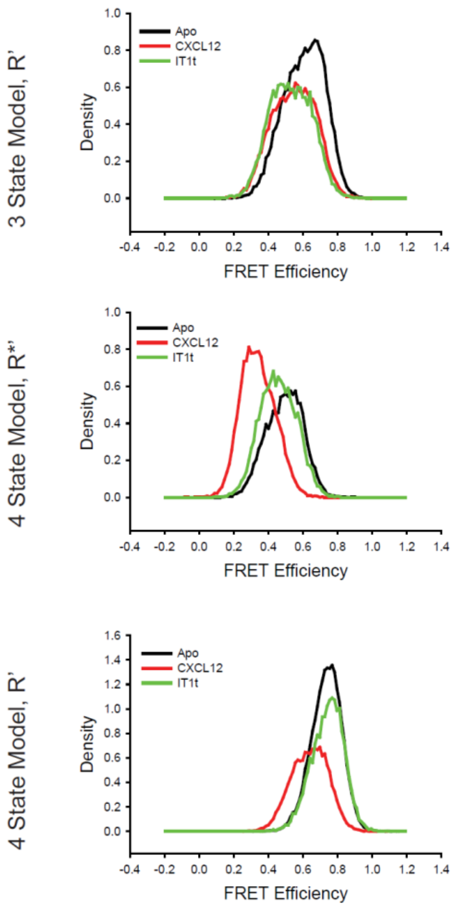


**Fig. S4.** Comparison of the CXCR4 intermediate states for the conditions detailed in Fig. S3 using three-state and four-state models revealed over fitting artifacts using four states. Top) With the 3-state model, the R' states for apo-CXCR4 and for CXCL12- and IT1t-bound receptor overlapped well with similar apparent FRET values across all of the tested conditions. In the case of the four-state model, the R*' (Middle) and R' (Bottom) states were substantially different across the ligand treatments. In particular, the R*' state with CXCL12 treatment appears to arise from a splitting of the R* conformation, indicating that the model was overfitting the data.

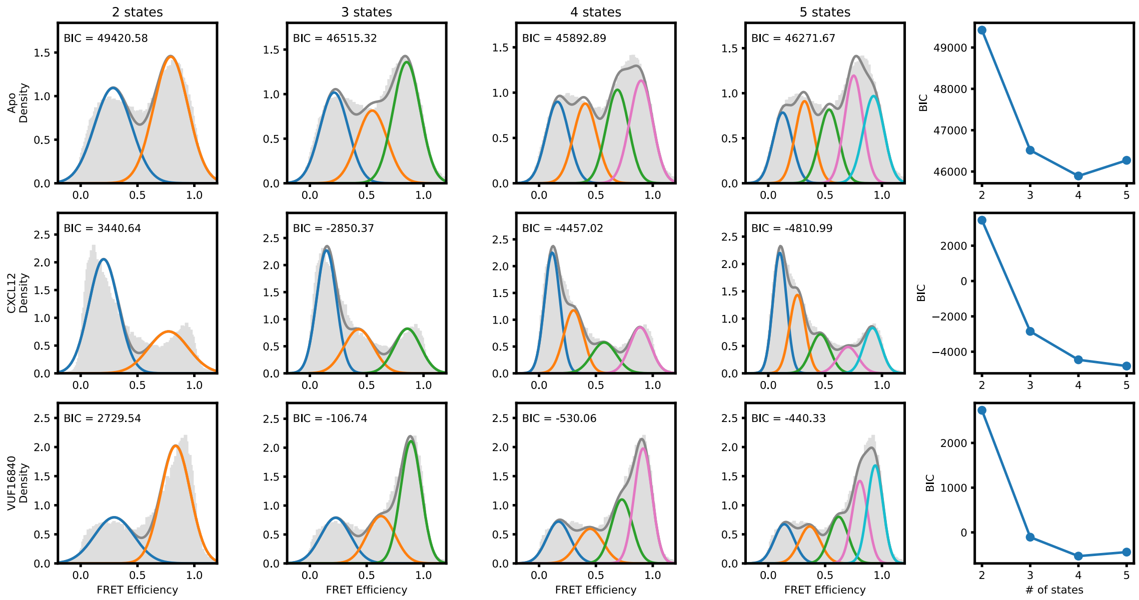


**Fig. S5.** Evaluation of the number of discrete FRET states present in ACKR3 under various conditions. SmFRET data for ACKR3 recorded in the apo-state (first row), and in the presence of agonist CXCL12$_{WT}$ (second row) and inverse agonist VUF16840 (third row) were globally analyzed by Hidden Markov analysis assuming the presence of two, three, four, or five FRET states. For each condition, the overall apparent FRET efficiency histogram is shown in gray. Analysis was performed as described in Fig. S3. The results indicate that four FRET states are required to reproduce the experimental distributions.

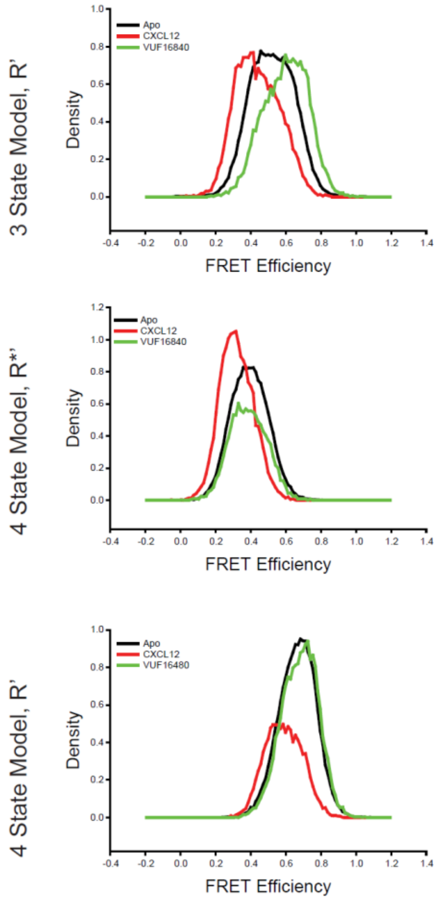


**Fig. S6.** Comparison of intermediate state FRET histograms of ACKR3 from the three- and four-state models shown in Fig. S5. Top) The R' state histogram from the three-state model for apo-ACKR3, and $CXCL12_{WT}$- and VUF16840-treated ACKR3 showed a spread across intermediate FRET values. The R' distribution of the apo-receptor was directly between the R' states for the agonist and inverse agonist samples. Splitting the single intermediate into two separate active-like (R*', Middle) and inactive-like (R', Bottom) with a 4-state model showed consistent state assignments across the intermediates, which indicated that the same conformations were being observed in each treatment. Therefore, four states were required to fully capture the observed conformational landscape of ACKR3.

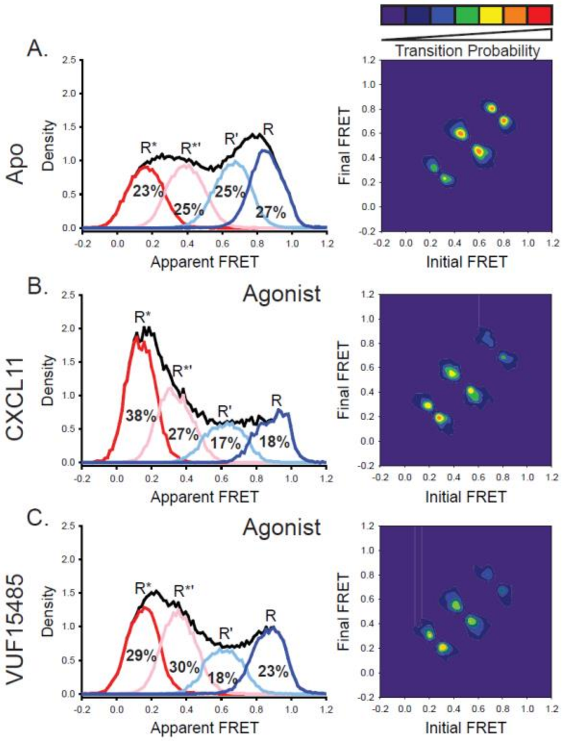


**Fig. S7.** The natural and engineered agonists CXCL11 and VUF15485, respectively, both promoted low-FRET, active-like ACKR3 conformations. A) Repeated results of apo-ACKR3 from Fig. 2B for comparison of the apparent FRET envelope and modeled states and the TDP. B) Treatment with CXCL11 leads to a predicted shift in population and transitions to low-FRET states. D) The small molecule agonist VUF15485 (1) also showed shifts of the population and transitions to low-FRET R* and R*' conformations. Results shown are the combined data sets of three independent experiments. The total apparent FRET envelopes for the samples are represented by black traces.

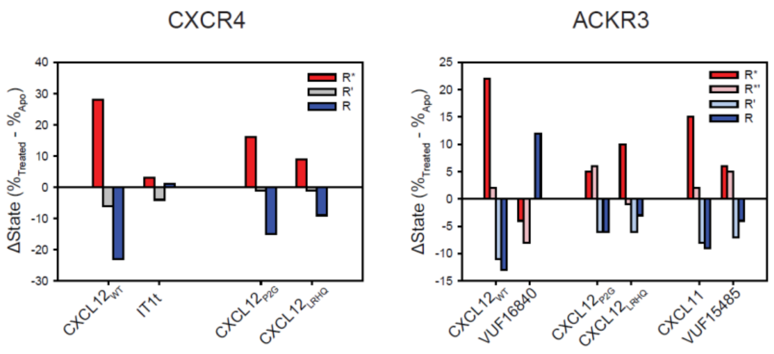


**Fig. S8.** Change in the population percentages of individual FRET states due to ligand treatment of CXCR4 and ACKR3. Data used for this analysis are presented as full apparent FRET distributions in Figs. 2, 3, and 4 and Fig. S7.

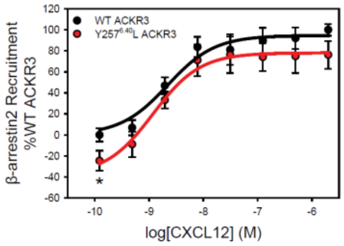


**Fig. S9.** The mutation Y257$^{6.40}$L reduces the constitutive activity of ACKR3. β-arrestin2 recruitment to ACKR3 was detected by BRET between the rluc-tagged receptor and GFP10-tagged arrestin. The data represent the mean of three independent experiments measured in triplicate with error bars reflecting the standard deviation. Statistical significance determined by unpaired t-test, *$P < 0.05$. The plot recapitulates similar data presented previously in Yen et al. 2022 (2) and is included here for reference.

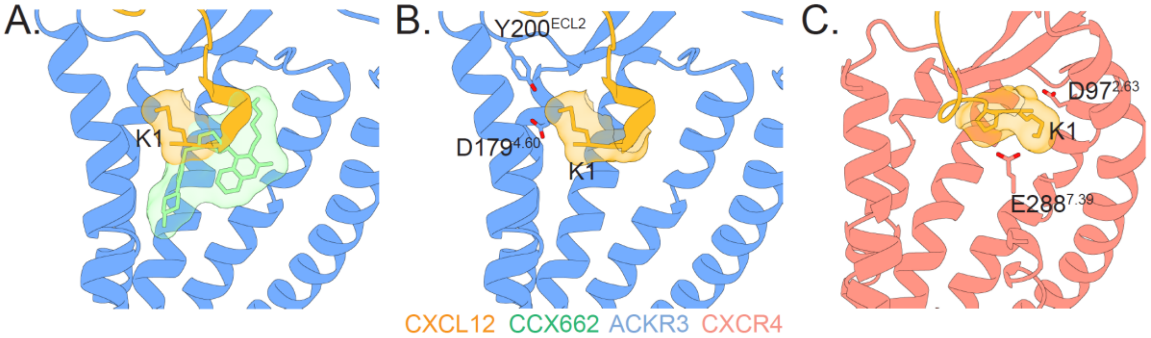


**Fig. S10.** ACKR3 is activated by many ligands, while CXCR4 requires specific interactions with the CXCL12 N-terminus for activation. A) Structures of ACKR3 (blue) bound to CXCL12 (gold, PDB ID 7SK3) or the small molecule agonist CCX662 (green, PDB ID 7SK8) reveal multiple modes of receptor activation (2). B) When CXCL12 binds ACKR3, the chemokine N-terminus is oriented to position CXCL12-K1 towards ACKR3-D179$^{4.60}$ and ACKR3-Y200$^{ECL2}$ near where the chemokine backbone enters the orthosteric binding pocket. C) In contrast, when bound to CXCR4, the N-terminus of CXCL12 is predicted to be in an extended conformation and positions CXCL12-K1 to interact with CXCR4-D97$^{2.63}$ and CXCR4-E288$^{7.39}$. The precise positioning of K1 appears critical, as mutation of P2 of CXCL12, which is surmised to orient K1, converts the agonist into an antagonist (3, 4). The model of CXCR4 was adapted from Ngo et al. 2020 (4) and Stephens et al. 2020 (5).
